# Rapidemic, a versatile and label-free DNAzyme-based platform for visual nucleic acid detection

**DOI:** 10.1101/2020.10.14.337808

**Authors:** Marijn van den Brink, Sebastian T. Tandar, Tim A. P. van den Akker, Sinisha Jovikj, Violette Defourt, Tom G. B. Langelaar, Tijn O. Delzenne, Kelly van Strien, Amber W. Schonk, Aukje J. A. M. Beers, Eugene Golov, Lucy J. Chong, Güniz Özer Bergman, Joey J. W. D. Meijdam, Marjolein E. Crooijmans, Dennis Claessen, Johannes H. de Winde

**Author notes:** Equal contribution authors (Marijn van den Brink, Sebastian T. Tandar).

## Abstract

In the last three decades, there have been recurring outbreaks of infectious diseases, brought to light with the recent outbreak of coronavirus disease 2019 (COVID-19). Attempts to effectively contain the spread of infectious diseases have been hampered by the lack of rapidly adaptable, accurate, and accessible point-of-care diagnostic testing. In this study, we present a novel design of a label-free DNAzyme-based detection method called Rapidemic. This assay combines recombinase polymerase amplification (RPA) with linear strand-displacement amplification (LSDA) and guanine-quadruplex (GQ) DNAzyme-catalysed colour-changing reaction. The colorimetry basis of the signal readout omits the need for extensive instrumentation. Moreover, the primer-based sequence detection of RPA gives Rapidemic a potential to be rapidly adapted to target a new sequence. As a proof of concept, we developed the assay to detect isolated genomic DNA of *Saccharomyces cerevisiae*. The use of low-pH buffers and the optimization of the dilution rates from each preceding reaction to the next showed to be successful strategies to enable visible detection with this method. These findings demonstrate for the first time that a label-free DNAzyme-based detection method can be coupled to RPA and LSDA for nucleic acid detection.

## Introduction

Outbreaks of infectious diseases cause major health challenges on a global scale. In the last 30 years, there have been recurring outbreaks of infectious diseases, such as Influenza, Dengue fever, Zika fever, Ebola, Yellow fever, cholera and Middle East respiratory syndrome (MERS)^1^. Recently, a novel severe acute respiratory syndrome (SARS), coronavirus disease 2019 (COVID-19), was discovered in China and spread worldwide, causing over 27,800,000 cases of infection and 900,000 deaths as of 10 September 2020^2^. The pandemic not only causes medical complications, but also has serious socio-economic and environmental impact^3,4^.

Attempts to effectively contain the spread of infectious diseases have failed due to the lack of rapid, accurate and accessible diagnostic testing^5,6^. Currently, diagnosis is routinely performed with polymerase chain reaction-based (PCR-based) tests^7,8^. PCR-based tests are accurate and fast, but the requirements for laboratory equipment and expert personnel increase the overall test time to over 24 hours^8^. The need for laboratory capacity is a restricting factor in population testing, especially in low-resource areas where lab testing is sparse^6,9^. Point-of-care rapid diagnostic tests exist, but these antigen-based tests often lack sufficient sensitivity^10–12^, and their clinical use for detecting COVID-19 is currently strongly discouraged by the World Health Organisation (WHO)^13^. In addition, the development of antigen-based tests takes longer than that of a PCR test^14^, while a rapid diagnostic response at the start of the outbreak is essential for successful outbreak containment^5,6^. To be prepared for potential future epidemic-causing pathogens such as listed in the WHO R&D Blueprint^15^ or unknown diseases collectively referred to as “Disease X”, there is an urgent need for innovative tests that can quickly be adapted to new pathogens.

Several nucleic acid amplification techniques have been developed as isothermal alternatives to PCR to avoid the need for thermocycling^16^. Examples of isothermal amplification methods are loop-mediated amplification (LAMP)^17^, nucleic acid sequences-based amplification (NASBA)^18^, rolling circle amplification (RCA)^19^, strand-displacement amplification (SDA)^20^, isothermal exponential amplification reaction (EXPAR)^21^ and recombinase polymerase amplification (RPA)^22^. Among other methods, RPA was found to be a promising method for versatile point-of-care applications due to its low reaction temperature and relatively simple primer design^22,23^. In the RPA reaction, a recombinase is used to assist the binding of two primers to the template DNA upon which a strand-displacement polymerase amplifies the target sequence. DNA and RNA targets of various organisms have been successfully amplified with RPA for rapid point-of-care diagnostics applications^24^. End point detection is usually performed using lateral flow strips or fluorescent probes. However, these methods may impair the simplicity of the test, as detection by lateral flow requires oligonucleotide labelling and immobilization of antibodies^25^, whereas fluorescent detection may require a specialized instrument to read the output signal. In addition, the colloidal gold and antibodies often used in lateral flow strips increase the cost of the test substantially^26^. Therefore, new simple and low-cost detection methods that can be coupled to isothermal amplification reactions would be valuable alternatives to the current techniques.

There has been increasing interest in the use of DNA enzymes (DNAzymes) with peroxidase-mimicking activity for molecular diagnostics^27,28^. DNAzymes are suitable for point-of-care testing due to their good chemical and thermal stability as compared to protein enzymes. They are amenable to various amplification methods, which can be employed to improve the sensitivity of the assays^27^. DNAzyme-mediated oxidation coupled to isothermal amplification methods has been described before^29,30^. Li et al. (2019) integrated a G3 DNAzyme with EXPAR for sensitive nucleic acid detection^29^. However, their mechanism of amplification severely limits the choice for a target sequence. Alternatively, Wang et al. (2019) exploited asymmetric amplification to couple RPA with DNAzyme-catalysed detection^30^. The asymmetric RPA resulted in the amplification of DNAzyme sequences that contained a large strand of bases adjacent to their 5’ end, which may possibly hamper with the peroxidase-mimicking activity of the DNAzyme. Moreover, asymmetric amplification methods such as asymmetric RPA often suffers from low amplification efficiency, case-specific optimization requirement, and generation of non-specific product(s)^31,32^.

We here describe Rapidemic, a simple label-free DNAzyme-based detection method for nucleic acid detection. This assay combines RPA with linear strand-displacement amplification (LSDA)^33^ and oxidation catalysed by a guanine-quadruplex (GQ) DNAzyme to yield a colour change that is visible to the naked eye^34–36^. As a proof-of-concept for this principle, we report the initial validation of this innovative detection method with genomic DNA of *Saccharomyces cerevisiae* (*S. cerevisiae*). Thereby, we demonstrate for the first time that a label-free DNAzyme-based detection method can be coupled to RPA and LSDA for efficient detection of nucleic acids.

## Materials and methods

### Genetic sequences, enzymes, and reagents

Primers and DNAzymes were purchased from Sigma Aldrich. Purified *S. cerevisiae* BY4741 genomic DNA was kindly obtained from Paul van Heusden (Leiden, Netherlands). Synthetic oligo DNA as LSDA templates were purchased from BaseClear (Leiden, Netherlands). Nicking endonucleases, including Nt.BstNBI, Nt.AlwI and Nt.BsmAI, and Bst 2.0 DNA Polymerase were acquired from New England Biolabs (NEB). Chemical reagents and dNTP Mix were purchased from Merck.

### RPA reaction

RPA was performed using the TwistAmp Basic RPA kit (TwistDx). RPA reaction mix was assembled by combining 29.5 μL TwistAmp Basic Rehydration buffer, 0.5 μM of each primer, 5 μL template and water in a total volume of 47.5 μL. Primers were designed using PrimedRPA^37^. 5’-overhangs included in reverse primers were added manually to the primers provided in PrimedRPA output. The TwistAmp Basic pellet was rehydrated by adding 47.5 μL RPA reaction mix and was incubated for 5 minutes at room temperature until completely dissolved. The reaction mix was mixed by pipetting and divided into 19 μL aliquots. To start the reaction, 1 μL of MgOAc (provided with TwistAmp Basic kit) was added. The reaction was incubated at 42 °C for 20 minutes. The reaction was terminated by PCR clean-up or by quenching the reaction on ice. After a routine clean-up using a PCR clean-up kit (QIAGEN), amplification was confirmed by agarose gel electrophoresis using a 2% agarose gel and a GeneRuler 100 bp DNA Ladder.

### LSDA reaction

The LSDA reaction mix was assembled on ice and contained 1 mM dNTP Mix, 0.5x NEBuffer 3.1, 0.5x Isothermal Amplification Buffer, 0.4 U Bst 2.0 DNA Polymerase, 3.2 U nickase, 1 μL RPA product and nuclease-free water in a total volume of 10 μL. The reaction mix was prepared in bulk and divided into 10-μL aliquots. The LSDA reaction mix was incubated at 55 °C for 40 minutes. Reactions were quenched by placing the tubes on ice.

### LSDA reaction for initial characterization of nickases

The LSDA reactions for the initial characterization of nickases were performed with synthetic templates (1.6 x 10^10^ copies/μL) to eliminate noise from the RPA reaction. Double Bst 2.0 Polymerase and nickase concentrations were used to accommodate signal production from the low template concentration. The LSDA reaction mix was assembled on ice and contained 1 mM dNTP Mix, 0.5x NEBuffer 3.1, 0.5x Isothermal Amplification Buffer, 0.8 U Bst 2.0 DNA Polymerase, 6.4 U nickase, 4 μL template and nuclease-free water in a total volume of 10 μL. Reaction mix was prepared in bulk and divided into 10-μL aliquots. The LSDA reaction mix was incubated for 40 minutes at 55 °C for Nt.BstNBI and 50 °C for Nt.AlwI and Nt.BsmAI. Reactions were quenched by placing the tubes on ice.

### GQ-catalysed TMB oxidation reaction

0.1 M phosphate citrate buffer (PCB, pH 3.8) was prepared by combining 7.1 mL of 0.2 M Na2HPO4.7H2O (MW 268.07 g/mol) and 12.9 mL of 0.1 M citric acid. The pH of the buffer was adjusted to 3.8 with HCl.

The wells of a 96-well plate were filled with 1.25 μL of LSDA product, 8.25 μL of 0.1 M PCB and 40 μL hemin/KCl solution, containing 1.165 mg/mL KCl, 2.5 μM hemin and 34 μL 0.1 M PCB. LSDA product and 8.25 μL of 0.1 M PCB were replaced with 10 μL 1 μM EAD2+3’A or 0.1 M PCB for positive and negative controls, respectively. To start the reaction, 50 μL of freshly prepared TMB/H2O2 start solution was added, containing 0.12 mg/mL TMB, 0.09% (w/w) H2O2 and 49.35 μL 0.1 M PCB. The addition of this start solution would result to a final TMB concentration of 0.06 mg/mL and a final H2O2 concentration of 0.045% (w/w). To stop the reaction, 50 μL 0.5 M sulfuric acid was added. The reaction was performed at 20 °C.

The solution’s absorbance at 650, 450, and 370 nm was measured every 40 seconds for 30 minutes with the Spark™ 10M multimode microplate reader. After the stop solution was added, the absorbance at 450 nm was re-measured.

### Real-time RPA reaction

Real-time RPA was performed by adding 2.5 μL of 20x SYBR green dye to a 50 μL RPA mixture. Before initiation of the RPA reaction with MgOAc solution, the whole RPA mixture was transferred to a 96-well RT-PCR plate. Signal measurement was performed using BioRad CFX96 Touch Real-Time PCR Detection system. Temperature was kept constant at 42 °C throughout the measurement. SYBR green signal readout was measured every 30 seconds.

## Results

### Rapidemic detection mechanism

The Rapidemic detection method relies on three main reactions: RPA, LSDA, and GQ-hemin DNAzyme-catalysed TMB oxidation. During RPA, the target sequence is amplified and extended with a short segment of nucleotides due to primer overhang (**Fig. 1a**). This segment contains the reverse complementary sequences of a recognition site for a nicking endonuclease (nickase) and a GQ sequence. The nickase makes a single-stranded cut near its recognition site in the amplification product, after which a strand-displacement polymerase elongates the 3’ end at the nickase cut site and thereby displaces the single-stranded GQ sequence, so that it is released from the double-stranded DNA molecule. The single-stranded GQ sequence then forms a three-dimensional structure with peroxidase activity upon binding to potassium ions and hemin (**Fig. 1b**). Finally, the DNAzyme catalyses the oxidation of 3,3’,5,5’-tetramethylbenzidine sulphate (TMB) in the presence of hydrogen peroxide (H2O2) to produce a colour change (**Fig. 1c**). As a proof-of-concept for this principle, we here report the initial validation of this detection method with genomic DNA of *S. cerevisiae*.

**Fig. 1.**
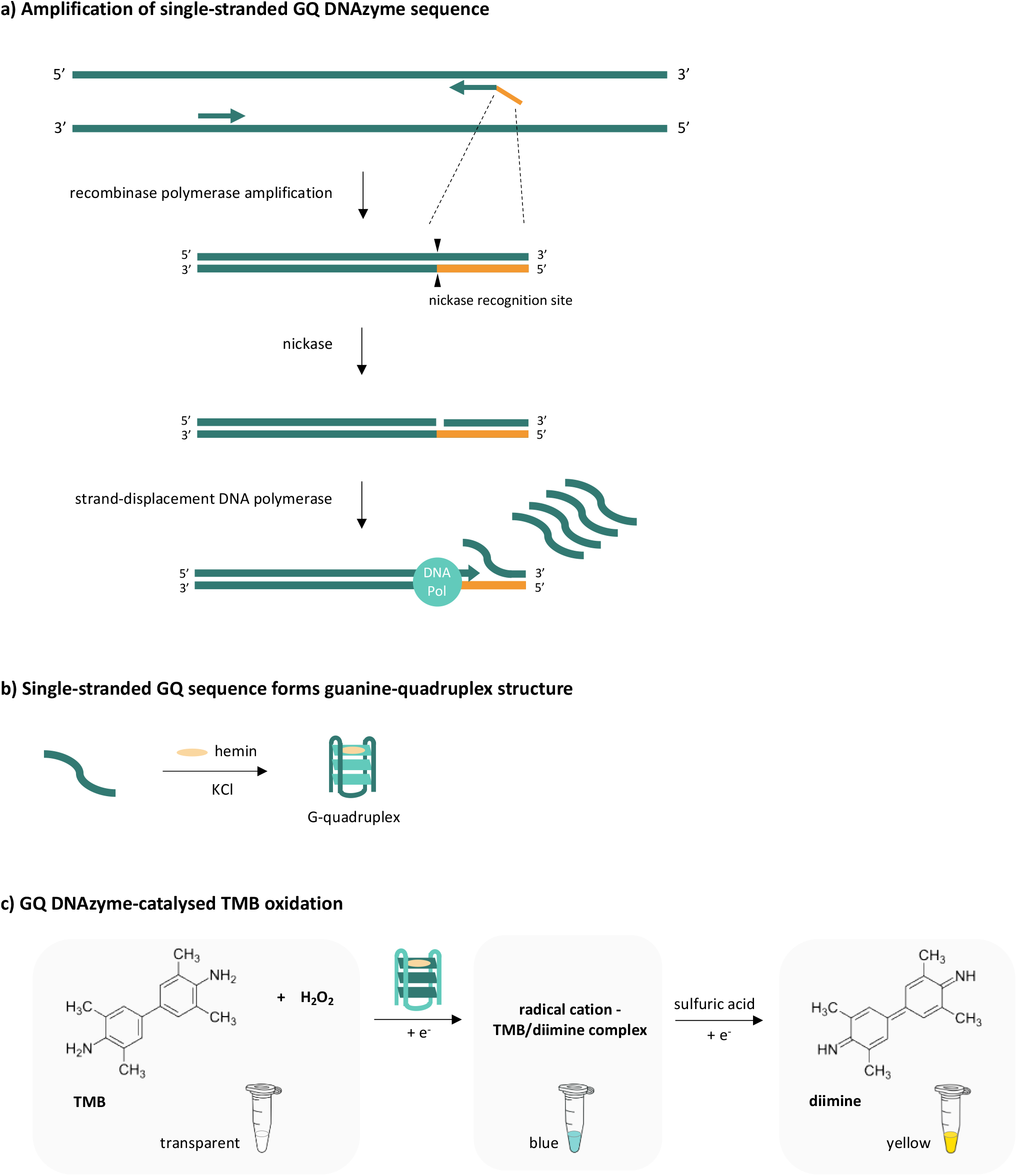
Mechanism of the DNAzyme-based label-free detection method Rapidemic.

### Confirmation of target amplification using recombinase polymerase amplification

Our method employed an initial target amplification step. This step is essential to selectively amplify the positive signal from the targeted sequence(s) present in the sample. Initial target amplification was performed using the TwistAmp Basic RPA kit (TwistDx). The primer pairs used in this experiment were designed to target a 191 bp sequence from the genome of *S. cerevisiae*. In addition to the target-specific sequences, the reverse primer contained a 27 bp overhang sequence at its 5’ - end. This primer overhang encoded for the reverse complementary sequences of a recognition site for a nickase and the EAD2+3’A DNAzyme, separated by a short spacer (less than 10 bp). Here we evaluated the primer pairs’ ability to detect the presence of its target sequence by using *S. cerevisiae* BY4741 genome as target sample (**Table S1**). As can be seen from **Fig. 2a**, the inclusion of 5’-overhang sequence did not interfere with RPA’s ability to detect its target sequence.

**Fig. 2.**
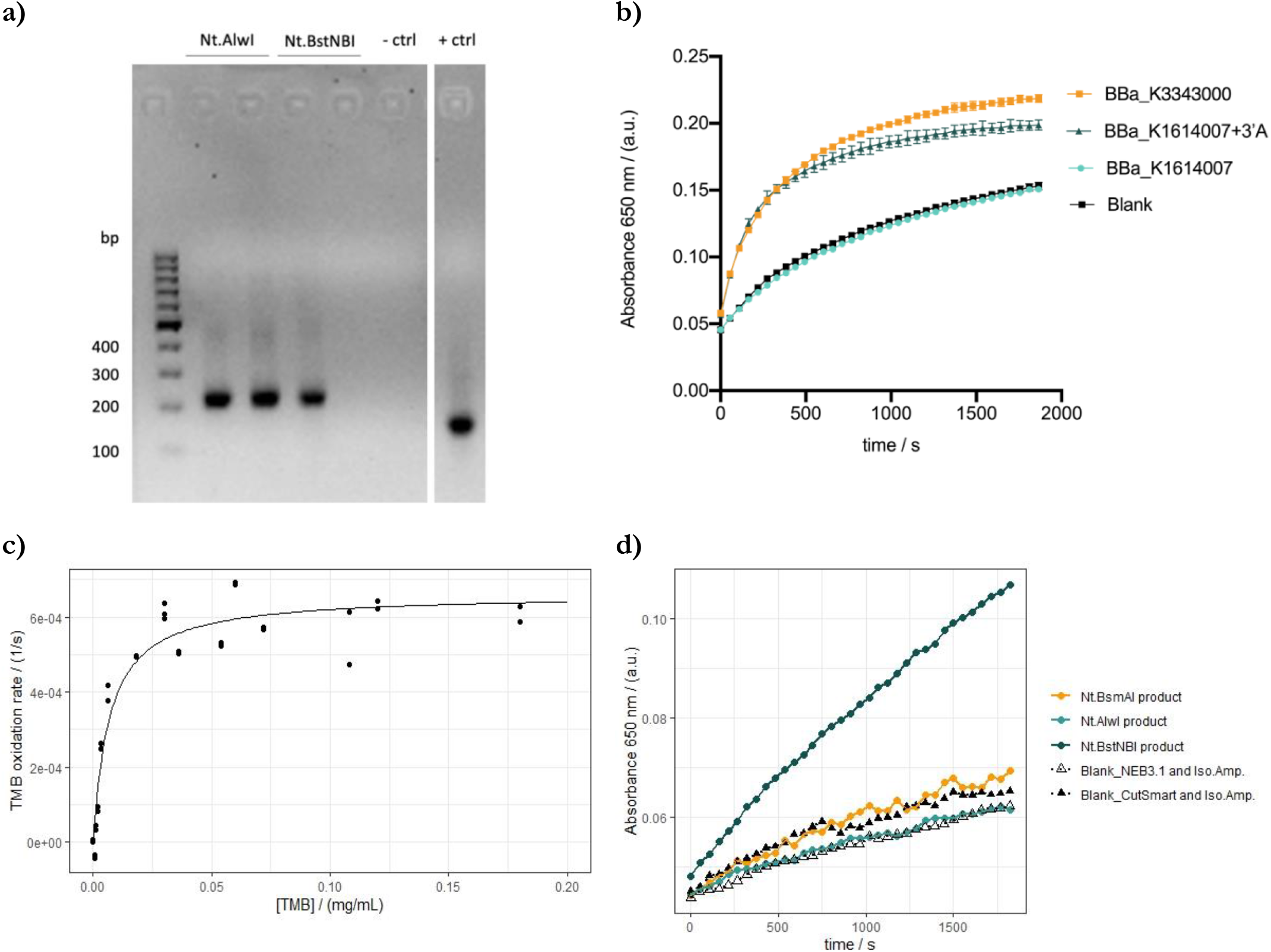
Confirmation of RPA, GQ-catalysed TMB oxidation and LSDA reactions. **a) RPA with SC1 forward and reverse primers containing Nt.AlwI or Nt.BstNBI recognition site**. Each reaction was performed using genomic *S. cerevisiae* BY4741 DNA as a target sequence. The target sequence is absent in the negative control. The positive control was performed with the primers and target templates provided by the TwistAmp Basic kit (TwistDx). **b) Average TMB oxidation rate of different GQ DNAzyme sequences.** Here, the catalytic activities of EAD2+3’A, BBa_K1614007, and BBa_K1614007+3’A GQ DNAzymes (Table S2) in pH 6.0 phosphate buffer were compared. The concentration of DNAzyme was kept at 1 μM in all measurements. EAD2+3’A was identified as the most potent DNAzyme in catalysing TMB oxidation. The error bars represented the standard deviation of three replicates. **c) TMB oxidation by EAD2+3’A DNAzyme.** Kinetic parameters of the DNAzyme were measured based on its ability to catalyse TMB oxidation reaction at different TMB concentrations. In the figure, data points represent initial oxidation rate measured from each experimental replicate while the curve represents model-predicted kinetics at each given TMB concentration. Rate kinetics of EAD2+3’A were assumed to follow Michaelis-Menten kinetics. The initial rate of each individual replicates was calculated based on differences in oxidized TMB absorbance value at 650 nm for the first minute of the observation. The resulting model could sufficiently describe each individual replicate measurement (Fig. S1). **d) GQ DNAzyme production using LSDA reaction.** LSDA was performed using different nickases. Here we identified Nt.BstNBI as the most potent GQ-producer. Synthetic (pure) oligonucleotides (Table S3) containing the targeted *S. cerevisiae* genome, nickase site and EAD2+3’A DNAzyme sequences were used as template for the LSDA reaction. The figure shows the average of two replicates.

### Confirmation of GQ-catalysed TMB oxidation

Guanine-quadruplex (GQ) sequences were known to have peroxidase-like activity when associated with hemin^36^. This allowed GQ-hemin complexes to be used as a catalyst for peroxidation reactions as an alternative to horseradish peroxidases^36,38^. Due to their enzyme-like ability, GQ-hemin complexes are often referred to as GQ DNAzymes. Just like horseradish peroxidases, GQ-hemin complexes can catalyse the colour-changing oxidation reaction of 3, 3’, 5, 5’-tetramethylbenzedine (TMB) in the presence of peroxide. However, different GQ sequences may harbour different enzymatic capabilities. Our preliminary analysis identified EAD2+3’A as the most potent GQ DNAzyme (**Fig. 2b**). EAD2+3’A was thus selected as final signal reporter.

Characterization of EAD2+3’A catalytic activity was performed based on the DNAzyme’s ability to oxidize TMB in the presence of excess amount of peroxide and different concentrations of TMB. All reactions were performed in a pH 6.0 phosphate buffer. The rate of TMB oxidation was then monitored based on oxidized TMB’s absorbance at 650 nm wavelength (molar absorption coefficient is 39,000 M^−1^ cm^−1^)^39^. Kinetic parameters of the DNAzyme were described using the Michaelis-Menten equation (**Eq. 1**), with *k_cat_* and *K_m_* representing the molar maximum TMB conversion rate and TMB affinity of EAD2+3’A, respectively. The values of *k_cat_* and *K_m_* were calculated using multiple regression analysis by simultaneously minimizing the error of all measurement-simulation pairs. The approximated kinetic parameters of EAD2+3’A against TMB are shown in **Fig. 2c, Fig. S1**, and **Table 1**.

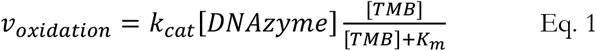

**Table 1.**
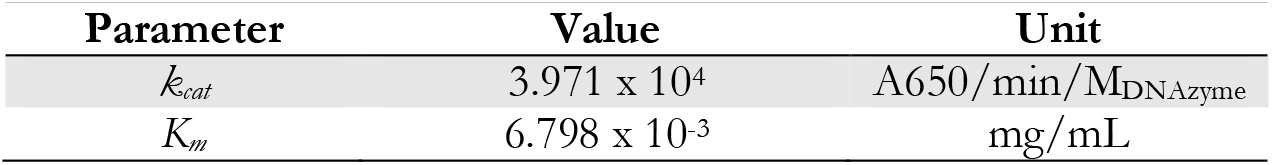
Kinetic parameters of EAD2+3’A in TMB oxidation reactions.

The *K_m_* value of the DNAzyme suggested that a TMB concentration of 0.06 mg/mL to be sufficient in order to reach 90% of the reaction’s maximum initial rate *k_cat_*. Further increase is TMB concentration may thus be not as significant in further increasing maximum reaction rate. Based on this result, a TMB concentration of 0.06 mg/mL was selected to be used in further assays and analyses.

### Confirmation of GQ DNAzyme generation with LSDA

In our sequence detection scheme, linear strand-displacement amplification (LSDA) acts as a connecting reaction to bridge initial signal generation by RPA and subsequent signal reporting by TMB oxidation. LSDA achieved this by producing GQ DNAzymes using purified RPA amplicons as templates. Here we confirmed the effectivity of LSDA in GQ sequence generation by using a synthetic (pure) template that resembles the RPA amplicon. The effectivity of Nt.BstNBI, Nt.Alwi, and Nt.BsmAI in mediating LSDA reactions was investigated. Two blank controls – which resembled the buffer conditions used in the three LSDA reactions – were included as standard of comparison. In the current reaction mix, we identified Nt.BstNBI as the most potent nickase for the production of GQ DNAzyme in an LSDA reaction (**Fig. 2d**). This result confirmed the effectivity of the LSDA reaction in producing single-stranded GQ DNAzyme as well as its role as a bridge between RPA and TMB oxidation.

### Dithiothreitol inhibits GQ oxidation

The feasibility of the detection method was then evaluated by passing the product of each preceding reactions to the next in a serial manner. Positive colorimetric signal from GQ-hemin-catalysed TMB oxidation could not be obtained using our original method and formulation (**Fig. 3a and b**). Interestingly, the signal could be obtained when RPA product was purified before being used in the subsequent LSDA reaction (**Fig. S2**). Thus, we suspected RPA mixture to contain a chemical(s) that may have inhibited LSDA and/or DNAzyme-catalysed TMB oxidation.

**Fig. 3.**
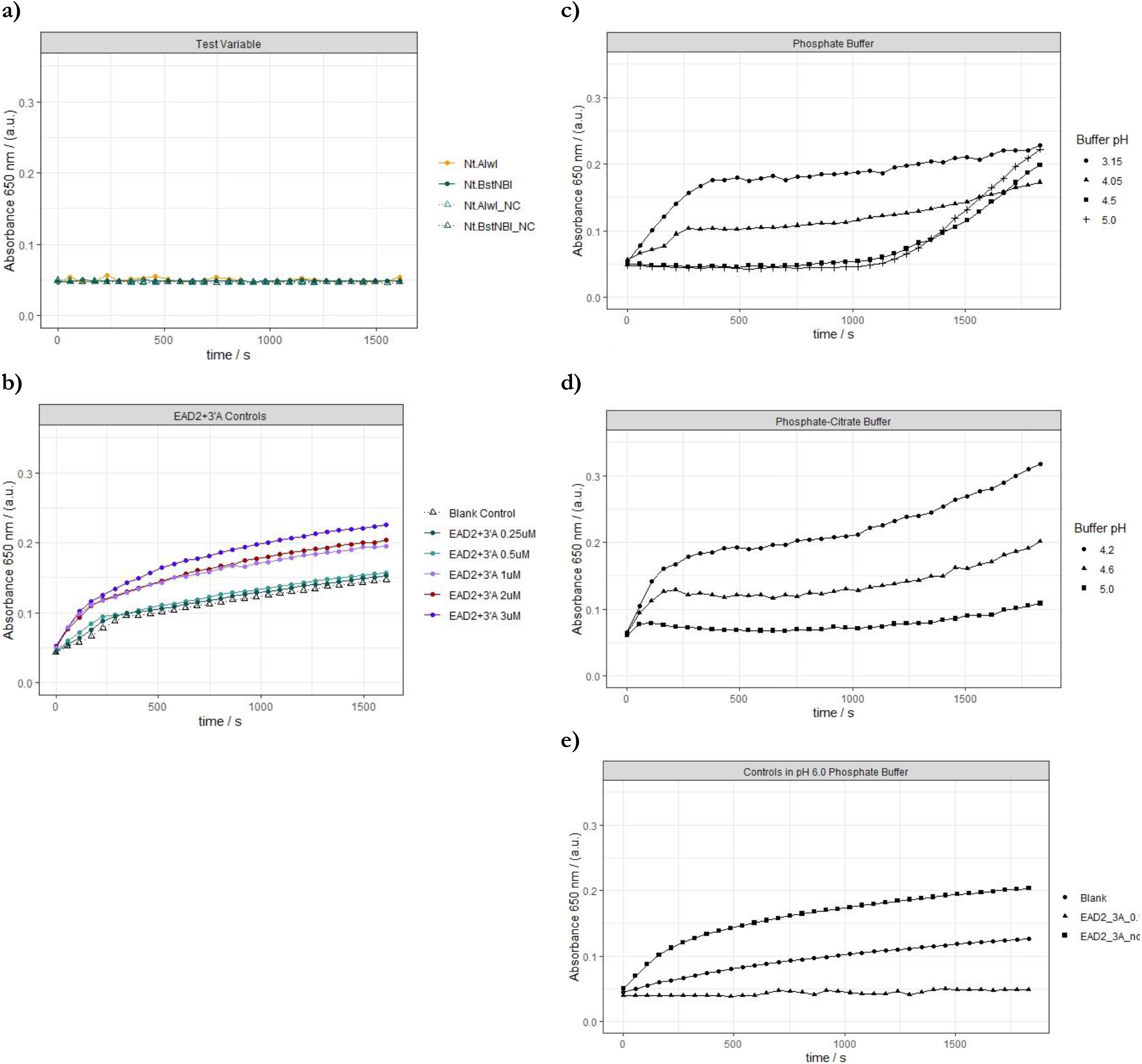
Effect of DTT on TMB oxidation reaction. **a and b) RPA mixture component prevented colour generation in TMB oxidation. a)** TMB oxidation of LSDA-treated unpurified RPA products could not generate observable colour change. The same was also observable from the negative controls (NC), which were performed in the absence of RPA target sequence. **b)** in a parallel measurement, colour generation could be observed when RPA mixture was not present in the final TMB oxidation mixture. Background colour generation from the blank control was also significantly higher than that observed from the test variables, suggesting the presence of a colour generation-inhibiting component in the RPA mixture. Data in a and b obtained from one replicate. **c, d and e) Acidic pH eliminates DTT reducing power.** Here, we investigated the suitability of acidic **c)** phosphate-citrate buffer and **d)** phosphate-citrate buffer for GQ DNAzyme-catalysed TMB oxidation reactions by comparing its performances to **e)** the original pH 6.0 phosphate buffer. In general, lower pH seemed to be favourable for signal production. Figures c, d and e show the average of three replicates.

The RPA mixture contained dithiothreitol (DTT), a strong reducing agent that is commonly mixed into protein formulations. The presence of DTT is known to improve protein stability by keeping the proteins’ monothiol groups in its reduced state^40^. Knowing its role as a reducing agent, we hypothesized that DTT carryover in TMB oxidation likely prevented the accumulation of oxidized TMB. Indeed, TMB oxidation with purified EAD2+3’A DNAzyme could not produce colour change in the presence of DTT (**Fig. S3**). Therefore, removal or inactivation of DTT is essential for our DNA detection scheme to work.

### DTT inactivation at low pH

The reducing power of DTT is known to be limited to more basic conditions^41^. Here, this characteristic of DTT was exploited as an alternative strategy to inactivate DTT during the DNAzyme-mediated TMB oxidation. Our original TMB oxidation formulation was performed using a pH 6.0 phosphate buffer. At this pH, we found that DTT hampers TMB oxidation (**Fig. S3**). Thus, we explored the possibility of DTT inactivation by performing DNAzyme-mediated TMB oxidation under different buffer conditions (**Fig. 3c, d and e**). Based on this comparison, phosphate-citrate buffer of pH 4.2 was identified to be the most suitable for TMB oxidation in the presence of DTT.

### Coupling RPA, LSDA, and TMB oxidation

Our attempt to serially couple RPA, LSDA, and TMB oxidation reaction in pH 4.2 phosphate-citrate buffer could not produce a significant colour signal. Although the low-pH buffer reduced DTT activity, the oxidation reaction was still inhibited by the presence of unpurified RPA mixture. **Fig. 4a and b** showed how the increasing amount of RPA mixture reduced the colour generation rate during the oxidation reaction. The pellet mixture was found to completely inhibit colour generation when diluted by less than 200-fold.

**Fig. 4.**
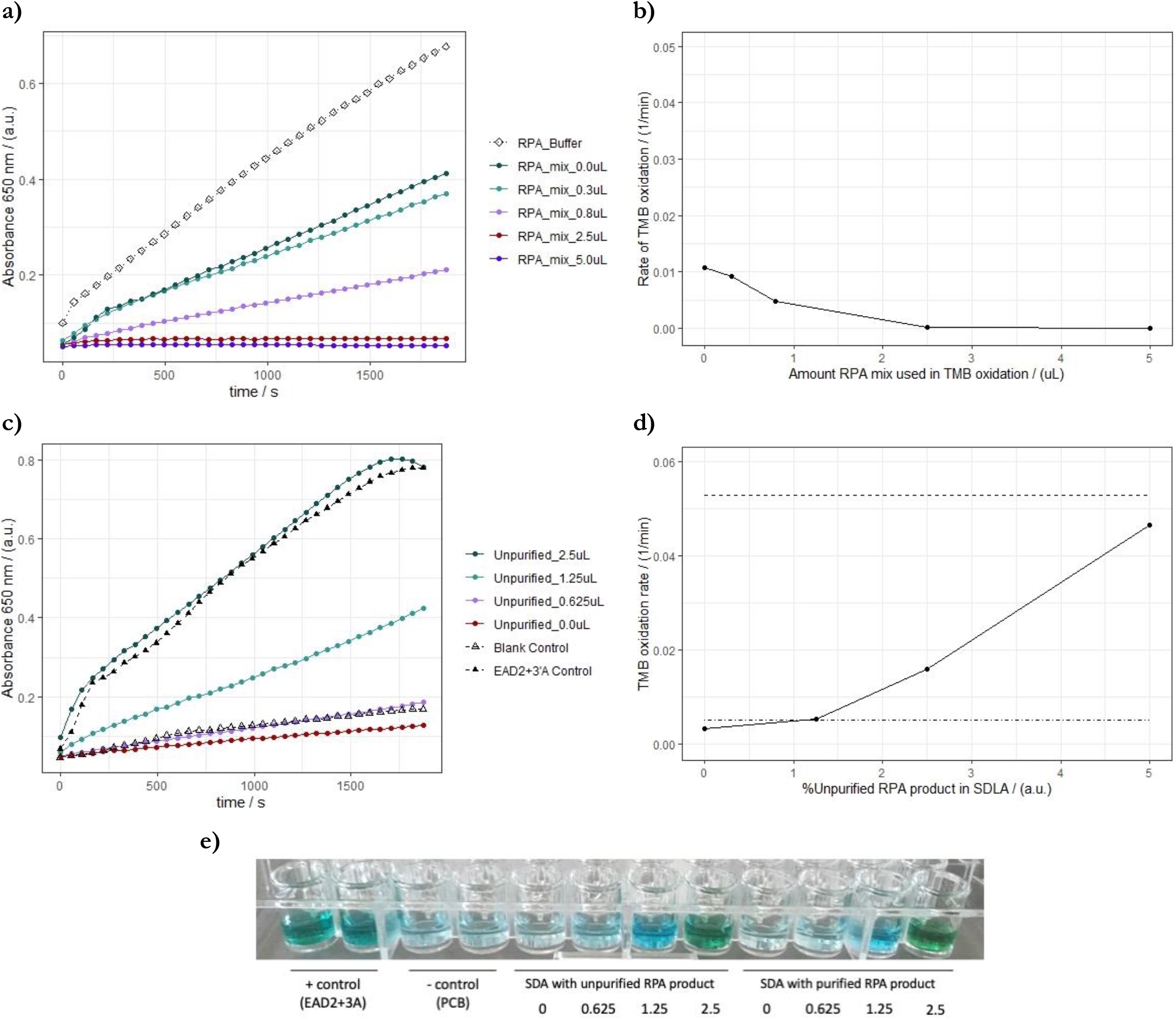
Dilution of RPA mixture is required for signal generation in TMB oxidation reaction. **a and b)** increasing the amounts of RPA mixture (not incubated, no template) was found to significantly reduce signal generation during the TMB oxidation reaction. In these reactions, 1 μM EAD2+3’A DNAzyme was used to catalyse the reaction. Figure a shows the average of two replicates. **c and d)** during TMB oxidation, a signal generation rate comparable to that achieved from 1 μM EAD2+3’A DNAzyme could be obtained by reducing the amount of unpurified RPA product that is passed on to the LSDA reaction to one-fourth of its amount in the original scheme. Within the observed RPA product concentration range, the initial TMB oxidation rate appeared to be correlated to the amount of unpurified RPA product used in LSDA. Figure c shows the average of two replicates. **e)** serial coupling of RPA, LSDA, and TMB oxidation using the updated dilution scheme and buffer could produce observable colour change.

To further attenuate the oxidation-inhibiting effect of the RPA mixture, we decided to reduce the amount of RPA and LSDA product used in each of its subsequent reactions in order to obtain a final RPA mixture dilution rate of at least 625-fold. Additionally, we decided to further increase the acidity of phosphate-citrate buffer used in the TMB oxidation reaction to a final pH of 3.8 to anticipate for potential pH increase when the solution was mixed with LSDA product, which had a slightly basic pH. Here we explored the effectivity of different dilution schemes from RPA to SDA and from SDA to TMB oxidation. Based on this analysis, a dilution of 10-fold during transfer from RPA to SDA and a subsequent dilution of 80-fold during transfer from SDA to TMB oxidation was found to be the most favourable dilution scheme (**Fig. 4c, d, and e**); the change in absorbance was similar to the positive control that did not contain any RPA or SDA reagents (no DTT).

Serial coupling of RPA, LSDA, and TMB oxidation was achieved by using the new dilution scheme and pH adjustment (**Fig. 4c and d**). The resulting colour signal was significantly higher compared to the blank controls. With the new dilution method and low-pH buffer, the TMB oxidation-inhibiting effects of compounds in the RPA and/or SDA reactions were diminished. Subsequently, this allowed successful coupling of RPA, SDA and TMB oxidation in a serial manner.

### DMSO reduced primer dimer formation during RPA and improved method accuracy

Our serial RPA-LSDA-TMB oxidation method could also generate colour signal even when the RPA reaction was performed in the absence of the targeted sequence (**Fig. 5a**). The false-positive signal was not observable when RPA mixture was not incubated. Similarly, the signal could not be generated when (pure) RPA primers were used as templates for the LSDA reaction (**Fig. S4**). This pinpointed the origin of the false signal to an RPA product. We suspected RPA product(s) of primer complexes to somehow generate a nickase site and a GQ sequence at the correct order, thus allowing LSDA to generate GQ DNAzyme from this side-product.

**Fig. 5.**
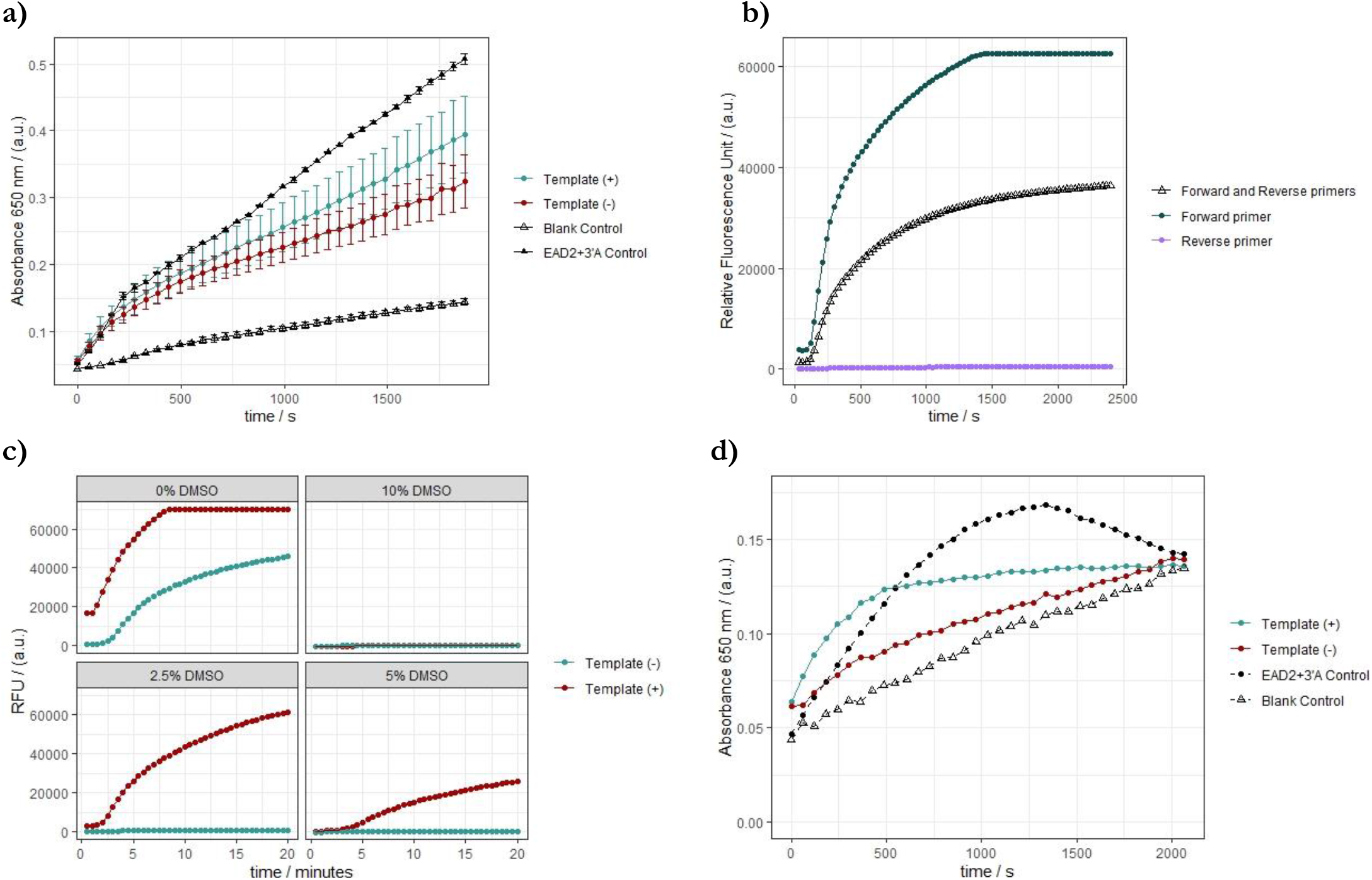
DMSO reduced primer dimer formation during RPA and improved method accuracy. **a and b) RPA amplification product in the absence of target template can produce a positive signal. a)** The average endpoint colorimetry readout of negative tests was approximately twice as high as was observed in the blank control. In this figure, error bars represented standard deviation of three (positive and negative tests) or two (blank and EAD2+3’A control) replicates. **b)** RT-RPA analysis confirmed the occurrence of background amplicon production in the absence of target template. Production of this background amplicon appeared to be dependent on the forward and not the reverse (tailed) primer. Data obtained from one replicate. **c) Addition of DMSO into RPA mixture could eliminate background amplicon generation.** Though effective, high concentrations of DMSO (5% v/v or higher) may also attenuate amplicon generation in the absence of target templates. Here we identified 2.5% DMSO to be the most suitable to distinguish the presence of template during the RPA reaction. Data obtained from one replicate. **d) 2.5% DMSO could significantly reduce false-positive signal from the overall sequence detection method.** Nevertheless, significant production of absorbance signal could still be observed 33.3% of the time. This may be due to residual production of false-positive-associated amplicons (Fig. S5). Average signals generated from positive- and negative-tests (n=6) are shown for comparison with the EAD2+3’A and the blank control. Individual measurements are shown in Fig. S6.

Real-time RPA analysis using SYBR Green dye captured the amplification of an unknown product in the absence of target template (**Fig. 5b**). This confirmed that our RPA primers alone could generate an amplicon. Interestingly, we also found that the non-tailed forward primer, and not the tailed reverse primer, could also generate a background amplicon on its own, even though the product may differ from that produced in the presence of both the forward and reverse primers.

As an attempt to reduce non-specific amplicon generation by primer complexes, dimethyl sulfoxide (DMSO) was added to the RPA reactions. DMSO is a polar aprotic solvent. Its ability to dissolve both polar and non-polar compounds allows it to disrupt secondary structure formation of DNA strands. By weakening the hydrogen bonds between primers, the presence of DMSO may prevent primer (self-) annealing^42,43^. Here we exploit this capability of DMSO to weaken the nonspecific primer-primer binding and complex formation (**Fig. 5c**). In this sense, DMSO acted much like a filter — eliminating background noise while allowing significant levels of true-positive signal to be further amplified. Here we identified 2.5% DMSO to be the most suitable for our RPA primer pair.

The inclusion of 2.5% DMSO in the RPA mixture was found to be effective in reducing the occurrence of false-positive signal from our sequence detection method. Significant attenuation of the true-positive signal was not observable, confirming the strategy’s effectivity to selectively filter out most false-positive amplicons (**Fig. 5d**). False-positive cases could nevertheless still be observable. We suspected this was due to the residual production of primer-associated amplicons. Although the current RPA condition could prevent false-positive signal production in RT-RPA, purified product from this reaction could still give a smear at approximately 100 bp region in agarose gel electrophoresis (**Fig. S5**). Nevertheless, the current method showed 33.3% false-positive and 33.3% false-negative rates based on production of absorbance signals higher than 0.15 (a.u.) within the given test period. This suggested the need for further improvements and technique optimization.

Interestingly, the maximum readout of true-positive signals was significantly lower compared to that observed in our previous experiments. This can be due to the interaction between DMSO and components of the LSDA and/or TMB oxidation reactions. Further analysis is required to pinpoint the cause of this signal attenuation and to further optimize our sequence detection technique.

## Discussion

In this study, we established the basic design of a label-free method for the detection of targeted nucleotide sequence. The method utilized a primer-based isothermal DNA amplification reaction known as RPA to detect the presence of a targeted nucleotide sequence. However, the use of RPA as a point-of-care detection method is rather limited by its inability to produce visible signal on its own. Here we showed how RPA can be coupled to a colorimetric reaction catalysed by a DNA-based enzyme (DNAzyme) known as guanine-quadruplex (GQ). When associated with hemin, GQ DNAzyme can serve as a peroxidase. This allows it to be used as an alternative to horseradish peroxidase in common oxidation-based colorimetry reactions, such as the oxidation of TMB and ABTS.

The DNAzyme’s essence as a DNA allows it to be produced alongside target amplification in RPA reactions. Our sequence detection scheme, the production of GQ DNAzyme was incorporated by encoding the reverse complement of GQ DNAzyme to the 5’-end of the RPA reverse primers used in our RPA reactions. As a result, this strategy allows GQ DNAzyme to be produced only when amplicon is produced during the RPA reaction.

GQ DNAzyme performs its catalytic activity as a single-stranded DNA. This creates a need to dispatch one strand of the co-amplified dsDNA GQ sequence from the rest of the RPA amplicon. In our sequence detection scheme, this role was fulfilled by linear strand displacement amplification (LSDA) reaction. LSDA relies on the activity of a nicking endonuclease to generate a single-stranded cut at the 5’-end of the GQ sequence as well as strand displacement capability of Bst 2.0 polymerase. Although the nickase Nt.BstNBI was used in most of our experiments, LSDA may also be performed by using other nickases, such as Nt.BspQI^33^.

As a whole, our sequence detection method incorporates three steps of amplification, reporter generation, and signal readout/measurement. The three reactions were RPA, LSDA and GQ-catalysed TMB oxidation, consecutively. The three reactions were connected in a serial manner such that a proportion of the product of each preceding reaction was used in each subsequent reaction.

Our first attempt to integrate the three reactions revealed how the presence of a reducing agent (commonly included in enzyme formulations) may have inhibitory effect in our colour-producing oxidation reaction (**Fig. 3a and b, Fig. S2**). In our experiments, DTT, which was inevitably present in the RPA and LSDA reaction mixes, was identified as the main reducing agent that prevented colour formation during subsequent TMB oxidation reactions (**Fig. S3**). The presence of DTT thus demanded a strategy to remove or inactivate DTT for our three-step detection scheme to function as intended.

Past attempts to couple RPA with oxidation-based colorimetry may have depended on buffer exchange procedures to remove DTT from the subsequent oxidation reactions. Though highly effective, these methods required additional measures to immobilize the reporting peroxidase while the previous solvent is washed away^44^. We circumvented the need of this additional complexity by implementing a dilution scheme that limits the DTT concentration in the final oxidation reaction. Knowing DTT’s significantly lower activity at acidic pH, we further adjusted our oxidation buffer pH to 3.8. By implementing the dilution scheme and pH adjustment of the oxidation reaction, we successfully coupled the three reactions in a serial manner (**Fig. 4c-e**).

Despite successful integration of the three reactions, the technique presented significant false-positive signals in the absence of target sequence. Further investigation revealed the production of primer-associated amplicons as the source of this false-positive signal. Using RT-RPA analyses, we found how the addition of 2.5% (v/v) DMSO in RPA reaction to effectively reduce the occurrence of false-positive signals (**Fig. 5c**).

The oxidation reactions that followed RPA with 2.5% DMSO could not yield sufficiently distinct true-positive signal when compared to the negative control and the blank controls. We hypothesize that this observation was the result of DMSO-GQ interaction. In this line, a study of Dhakal et al. (2013) reported that coexistence of parallel and non-parallel GQ species (1:1 ratio) was observed in the presence of crowded buffers with dehydrating cosolutes, such as 40% w/v DMSO^45^. The fact that complexes formed by hemin and antiparallel GQs have much lower peroxidase-mimicking activity than those formed by hemin and parallel GQs may explain the low readout signal that was observed when DMSO was included in the RPA reaction^46^. This revealed the need to further improve and optimize the presented sequence detection method to be used as a rapid diagnostic tool.

Despite the need for further optimization, the presented sequence detection method holds potential to be a rapid diagnostic tool. Its RPA-based detection allows it to be repurposed to target other DNA sequence by simply changing its primers, similar to the development of PCR-based tests. This allows rapid adaptation of the method to target newly emerging diseases. In comparison, development of antibody and/or antigen-based tests may require significantly more time and resources to be redeveloped to target a new disease^14^. Additionally, the reagent costs of the presented detection method are similar to commercial PCR-based tests (Supplementary Data). However, unlike PCR-based tests, our method does not require extensive instrumentation to give a signal readout. Altogether, the method presents a potential to be a rapidly adaptable point-of-care diagnostic tool.

Future research may focus on combining RPA and LSDA into a one-pot reaction. The development of a one-pot reaction may further hasten the reactions’ incubation period and reduce the reagent costs (dNTPs, buffers, polymerase). Moreover, this would greatly simplify the hardware design when applied in point-of-care diagnostics. It may be necessary to replace the commercial TwistDx RPA kit with a custom-made RPA mix to optimize its composition for one-pot RPA and LSDA. With such custom-made mix, the concentration of DTT could be reduced to increase the amount of RPA product allowed in the oxidation reaction and increase the method’s sensitivity. Altogether, the presented sequence detection scheme presented a promising potency as rapid point-of-care diagnostic tool that may contribute in future attempts to diagnose, and by doing so, monitor and contain the spread of an infectious disease.

## Supporting information

Supplemental Cost analysis

## Acknowledgements

This project was conducted as part of the International Genetically Engineered Machine (iGEM) Competition by team iGEM Leiden 2020. We thank Floor Stel, Jonah Anderson, Laurens ter Haar and Maarten Lubbers for support and encouragement during the project. We thank Paul van Heusden for providing us with *S. cerevisiae* genome.

## Contributions

Experiment was designed by S.T., S.J., M.B. and T.A, performed by S.T., M.B., T.A., V.D, T.L, T.D, A.S., K.S. and S.J. under the supervision of J.W., M.C., and D.C. Analysis and drafting was performed by M.B. and S.T. All authors contributed equally in project ideation and manuscript finalization.

