## Supplemental Figures S1-6 and Tables S1-3 for "Rapidemic, a versatile and label-free DNAzyme-based platform for visual nucleic acid detection"

### Supplementary Materials

**Table S1. RPA primers targeting *Saccharomyces cerevisiae***

| Primer | Sequence (5' to 3') |
|---|---|
| SC1_F | ACATTCTGTTTGGTAGTGAGTGATACTCTT |
| SC1_R_Nt.AlwI_EAD2+3'A | TCCCTCCCTCCCTCCCAGAGATGATCCTCTTATCGATAACGTTCCAATACGCTCAGT |
| SC1_R_Nt.BstNBI_EAD2+3'A | TCCCTCCCTCCCTCCCAGTCCAGACTCTCTTATCGATAACGTTCCAATACGCTCAGT |

**Table S2. GQ DNAzyme Sequences used in this Study**

| Name* | Sequence (5' to 3') |
|---|---|
| EAD2+3'A | CTGGGAGGGAGGGAGGGA |
| BBa_K1614007 | GGGTAGGGCGGGTTGGG |
| BBa_K1614007+3'A | GGGTAGGGCGGGTTGGGA |

* Alternative names reported in literature for BBa_K1614007 and BBa_K1614007 are Dz-00 and Dz-11, respectively[36].

**Table S3. Synthetic genomic sequences of *Saccharomyces cerevisiae* extended with a sequence containing a recognition site for a nickase and GQ DNAzyme EAD2+3'A**

| Name | Sequence (5' to 3') |
|---|---|
| Nt.AlwI / EAD2+3A | GTAATGTGAATTGCAGAATTCCGTGAATCATCGAATCTTTGAACGCACATTGCCCCCTTGGTATTCCAGGGGGCATGCCTGTTTGAGCGTCATTTCCTTCTCAAACATTCTGTTTGGTAGTGAGTGATACTCTTTGGAGTTAACTTGAAATTGCTGGCCTTTTCATTGGATGTTTTTTTTCCAAAGAGAGGTTTCTCTGCGTGCTTGGGATCATCTCTGGGAGGGAGGGAGGGA |
| Nt.BstNBI / EAD2+3A | GTAATGTGAATTGCAGAATTCCGTGAATCATCGAATCTTTGAACGCACATTGCCCCCTTGGTATTCCAGGGGGCATGCCTGTTTGAGCGTCATTTCCTTCTCAAACATTCTGTTTGGTAGTGAGTGATACTCTTTGGAGTTAACTTGAAATTGCTGGCCTTTTCATTGGATGTTTTTTTTCCAAAGAGAGGTTTCTCTGCGTGCTTGGAGTCTGGACTGGGAGGGAGGGAGGGA |
| Nt.BsmAI / EAD2+3A | GTAATGTGAATTGCAGAATTCCGTGAATCATCGAATCTTTGAACGCACATTGCCCCCTTGGTATTCCAGGGGGCATGCCTGTTTGAGCGTCATTTCCTTCTCAAACATTCTGTTTGGTAGTGAGTGATACTCTTTGGAGTTAACTTGAAATTGCTGGCCTTTTCATTGGATGTTTTTTTTCCAAAGAGAGGTTTCTCTGCGTGCTTGGTCTCGCTGGGAGGGAGGGAGGGA |

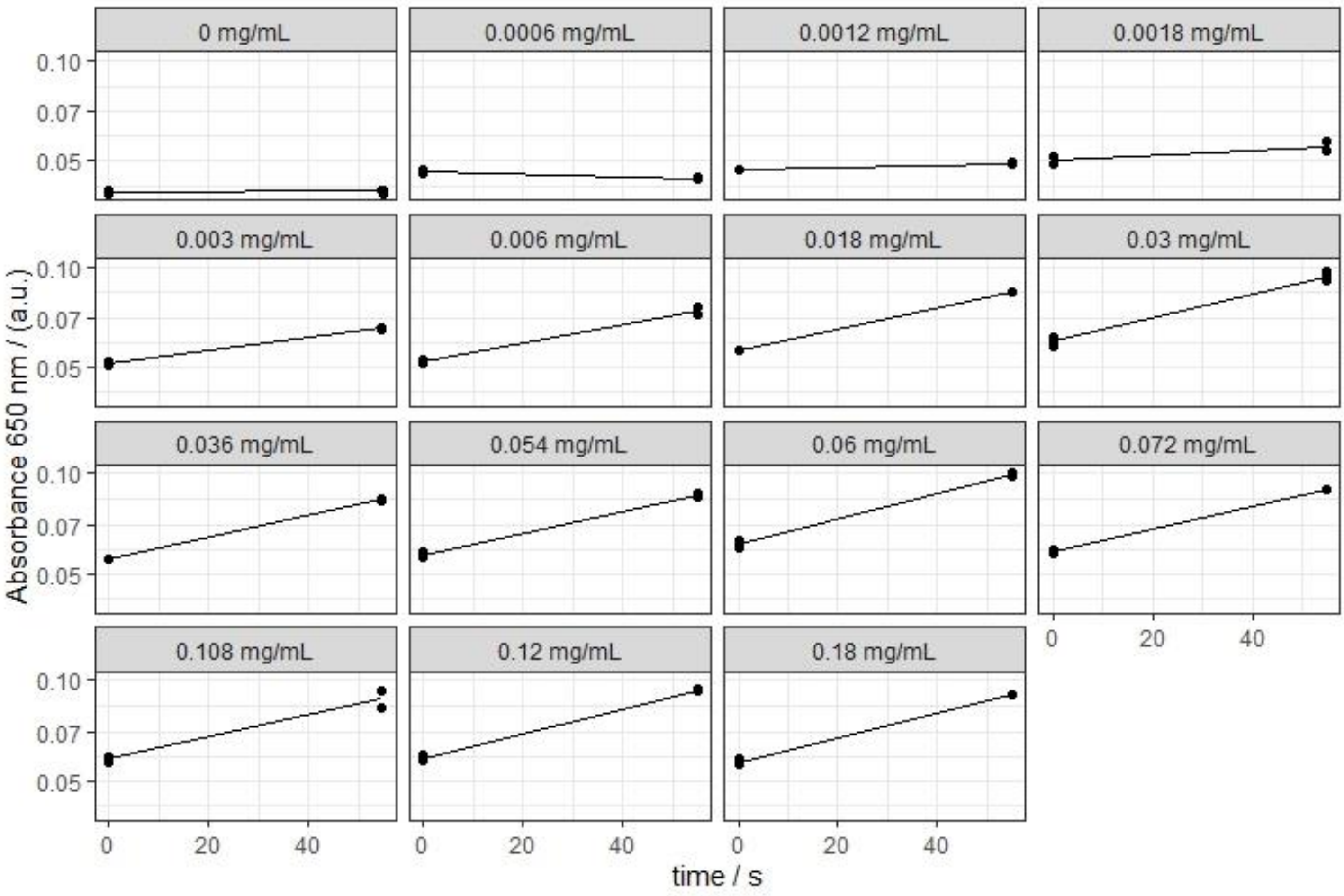


**Fig. S1 | Predicted TMB oxidation kinetics by EAD2+3'A could sufficiently describe reaction rates of each individual reactions.** The values above each figures represented TMB concentration used in each observation. Observations obtained from each experimental replicates were represented as data points. Reaction rate between each two observed time points were assumed to be constant and resulted to a linear increase in $A_{650}$ value. Lines shown in the figure represent the simulated increase in A650 value over the observation period at a given TMB concentration, as predicted by the fitted kinetic model (Fig. 2c), and by using the average of each measurement replicates at time point zero as starting point of the simulation.

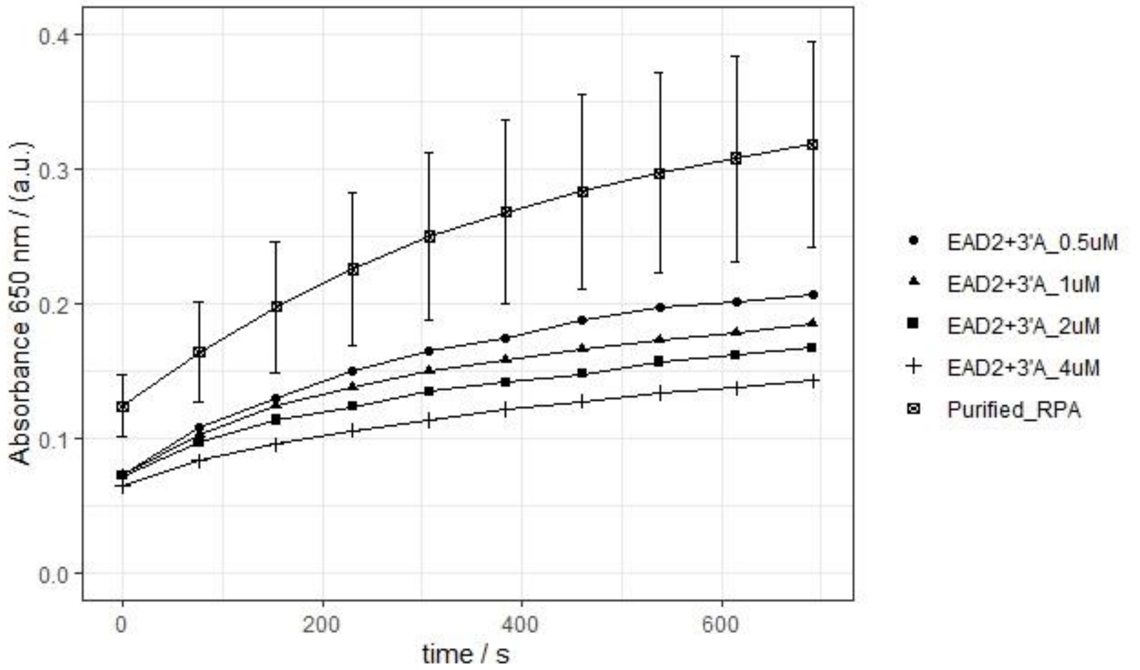


**Fig. S2 | TMB oxidation could be performed using LSDA product from purified RPA product.** TMB oxidation progressed at a significantly higher rate in the presence of LSDA product compared to when the reaction was spiked with pure 4 μM EAD2+3'A DNAzyme. This suggested LSDA's effectivity in producing EAD2+3'A to more than 4 μM within the 40-minutes incubation time. Moreover, this suggested the importance of RPA mixture removal in order to obtain TMB oxidation signal. The error bars represented the standard deviation of two replicates.

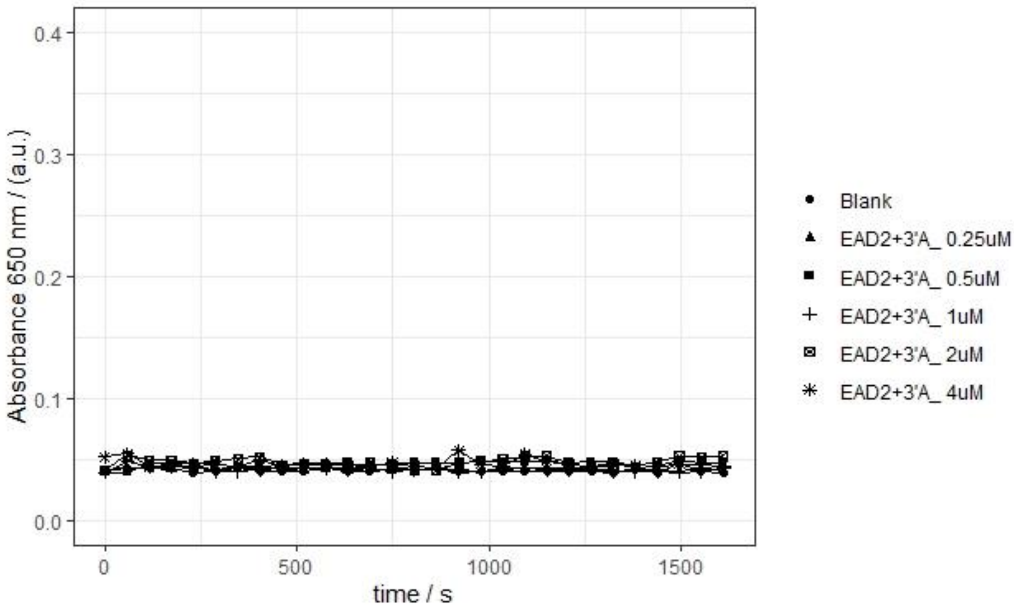


**Figure S3 | DTT inhibits TMB oxidation reaction by EAD2+3'A.** Absorbance signal at 650 nm of each spiked samples was not distinguishable from the background noise signal in the presence of 0.1 mM DTT. Figure shows the average of two replicates.

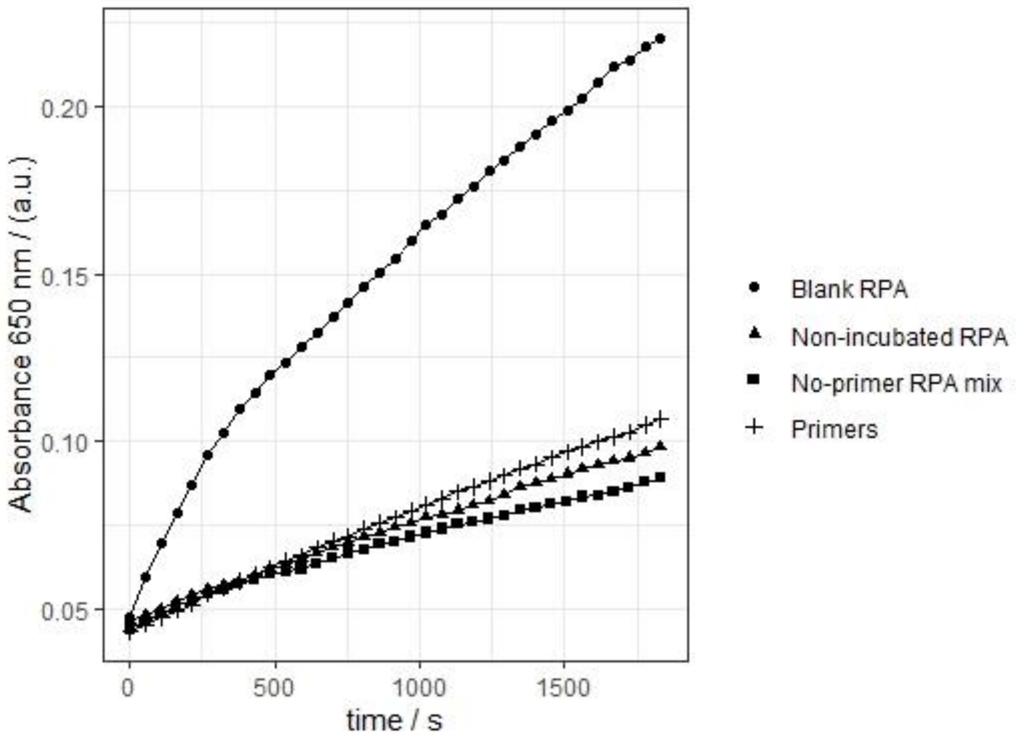


**Fig. S4 | False-positive signal originated from RPA side-product in the absence of target template DNA.** Without incubation, the same no-template RPA mix could not produce similarly high signal during TMB oxidation (triangles). The same is also true when either primer sets or RPA mixture was missing. Data obtained from one replicate.

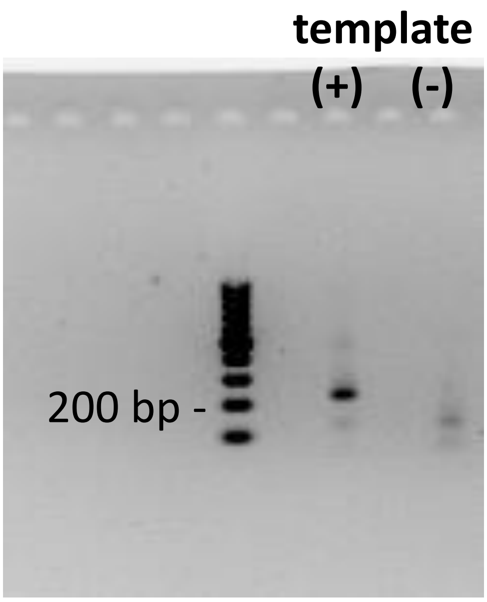


**Fig. S5 | Faint band at approximately 150 bp region could be observed from purified RPA product in the absence of target template and in the presence of 2.5% DMSO.** Although this band could only become visible at a higher resolution, this suggested residual production of false-positive-associated amplicons that may result to an increase in our method's false-positive rate.

**a)**

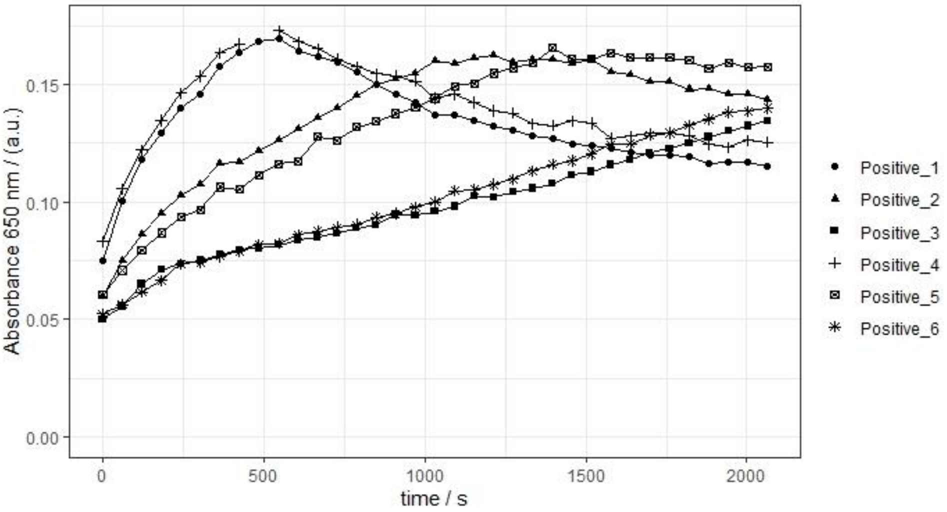


**b)**

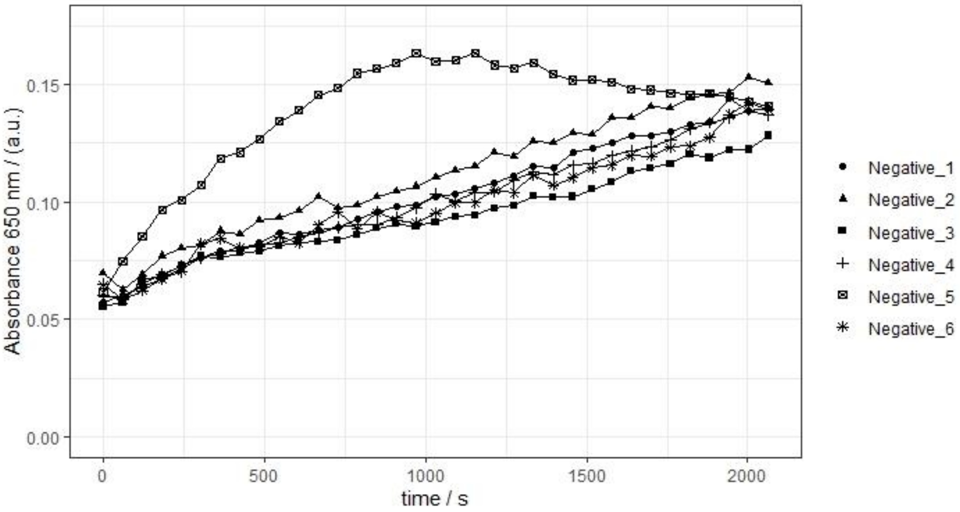


**Fig. S6 | 2.5% DMSO could significantly reduce false-positive signal from the overall sequence detection method. a)** Positive RPA reactions (with target sequence) and **b)** negative RPA reactions (without target sequence) were performed in presence of 2.5% DMSO and analyzed with LSDA and GQ-catalysed oxidation. Average signals generated from each positive- and negative-tests are shown in Fig. 5d for comparison with the EAD2+3'A and the blank control.
